# Oxidative stress and Atherosclerotic Plaque Progression: A plausible insight into the role of paraxonases and Ox-LDL

**DOI:** 10.1101/2025.10.28.685182

**Authors:** Divya Bhavani Ravi, Lakshmi Narasimhan Chakrapani, Thangarajeswari Mohan, Kalaiselvi Periandavan

**Author notes:** Correspondence: Dr. P. Kalaiselvi, Professor & Head; (P.K.).

## Abstract

The atherosclerotic heart disease is a complex disease associated with a plethora of dynamics contributing to its initiation and progression. Despite, diabetes and hypertension provide favorable conditions that provoke and aggravate atherosclerosis, it is undeniable that all of them do not have the tendency to develop plaques. This study was devised to identify the attributes that discriminate the diabetic and/or hypertensive ischemic patients from their corresponding non-ischemic population. Serum total antioxidant status, ox-LDL levels, Paraoxonase levels and its activity were assessed in the study groups. In order to evaluate the extent of the atherosclerotic disease progression, the aortic punch tissue samples from patients undergoing CABG were analyzed for the expression of ox-LDL and PON-2. Our results showed that their serum levels correlated well with the tissue expression. To ascertain the same with the progression of the disease, animal studies with rats fed with High Cholesterol Diet were carried out. Our findings suggests that ox-LDL and HDL-PON activity reflects the atherogenic events taking place in the arteries and further assessment of atherosclerotic risk in terms of ox-LDL and PON might be recommended after further validation.

## 1. Introduction

Cardiovascular diseases (CVD) endure to be a massive obstacle to the sustenance of human race and development contributing to a major percentage of mortality rates worldwide [1]. A close association is said to subsist between diabetes and cardiovascular diseases, the most prevailing major contributing cause towards the morbidity and mortality in patients with diabetes [2]. Patients with diabetes have a twofold risk of developing hypertension and patients with hypertension have a substantially higher incidence of developing diabetes [3]. It can be alleged that diabetes is greatly associated with the risk of cardiovascular complications, which is inflated in the presence of coexistent hypertension.

Ischemic Heart Diseases (IHD), a condition presented with narrowing of coronary arteries is caused by atherosclerosis compromising the blood and oxygen supply to the heart [4]. Even though atherosclerosis is a complex disease associated with a plethora of factors, the dysfunctional endothelium getting permeable to the entry of circulating lipoproteins (low-density lipoprotein, LDL) marks the initiation of the atherosclerotic events. With time, thus accumulating LDL undergo various changes especially oxidative modifications forming oxidized LDL (Ox-LDL) especially in conditions of diabetes where the ROS generation is exacerbated and therefore more oxidation of LDL leading to cardiovascular disease progression [6]. Besides, hypertension can be regarded as a major risk factor for this process as it results in increased stress of the vessel wall and might possibly augment the permeability of lipoproteins favoring the entrapment of lipids and lipoproteins within the arterial intimal walls [5]. Thus it can be alleged that the oxidative stress and its associated inflammation are critical aspects of atherosclerotic disease development and progression.

Paraoxonase (PON1) is an enzyme associated with high-density lipoprotein (HDL) and has a renowned role in serving as a cardio protective agent, as it guards both low-density lipoproteins (LDLs) and high-density lipoprotein (HDLs) against oxidative injure thereby curtailing the formation of ox-LDL [7]. Kotur‐ Stevuljević et al. [8] proposed that the decreased protective activity of HDL against peroxidation is attributed to the diminished activity of PON1 in the diabetic patients.

Lipid profile investigation along with the lipoprotein ratio assessment is regarded to be the regular laboratory investigation employed to assess the atherogenic risk intensity and to decide the medication strategy to either lower the LDL cholesterol /increase the HDL cholesterol. Although this assessment is quite easy and commonly employed, the efficacy of these lipoprotein remains controversial because, LDL does not stimulate any inflammatory response nor leads to foam cell formation unless it gets oxidatively modified (Ox-LDL) and the athero-protective efficacy of HDL does not rely only on its levels alone but also on its functionality in terms of its association with PON. Although Ox-LDL and PON have been already established as risk predictors of atherogenesis in different studies, their insinuation in the progression of atherosclerotic disease progression can be regarded as an important aspect towards arriving at risk prediction strategies.

## 2. Materials and Methods

### 2.1. Experimental Design 1 (Human Studies)

Blood samples were collected from control population attending the general outpatient ward and also, from cardiology and diabetology ward of the Government Kilpauk Medical College &Hospital (GKMC&H), Chennai after obtaining prior consent from them (IHEC Approval No: UM/IHEC/08-2013-I). Selection of individuals based on inclusion and exclusion criteria was mentioned in the supplementary file 1.

The Study group was devised into following: Group-I (Apparently healthy individuals); Group-II (Patients with a history of atherosclerotic risk factors alone and no apparent CVD and they are subdivided into IIa - Patients with the history of Hypertension, IIb - Patients with the history of Type 2 Diabetes, IIc - Patients with the history of Type 2 Diabetes and Hypertension); Group-III (Patients on admission for myocardial ischemia with a history of respective risk factors and they are subdivided into IIIa - Hypertension, IIIb - Type 2 Diabetes, IIIc - Type 2 Diabetes and Hypertension. All patients were on medication for their corresponding clinical conditions except for MI. Anthropometric measurement of the study populations are mentioned in the supplementary file 2 as Table 1. Biochemical parameters such as glucose, urea, Creatinine, total cholesterol, LDL, VLDL, TG and HDL measured among the experimental groups and results are presented in supplementary file 2.

### 2.2. Experimental Design 2 (Animal Studies)

Aged male albino rats of Wistar strain were obtained from Central animal facility, Taramani campus, University of Madras and maintained as per national guidelines and protocols, approved by the Institutional Animal Ethics Committee (IAEC No:02/08/2017). All experiments with animals were performed in compliance with the relevant laws and institutional guidelines. The animals were divided into four groups of six animals each. Group I-Control rats fed with normal rat feed for 45 days; Group II - Rats fed with High Cholesterol Diet (HCD) (4% cholesterol and 1% cholic acid) for 15 days [9]; Group III-Rats fed with High Cholesterol Diet (HCD) (4% cholesterol and 1% cholic acid) for 30 days; Group IV-Rats fd with High Cholesterol Diet (HCD) (4% cholesterol and 1% cholic acid) for 45 days. Biochemical parameters such as total cholesterol, LDL, VLDL, TG and HDL measured among the experimental groups and results are presented in supplementary file 2 as Table 2.

At the end of the experimental period, blood was collected from the rats, then the rats were anesthetized with ketamine (22 mg/kg b.wt, i/m) and aorta were excised immediately, immersed in ice-cold physiological saline and weighed. A 10% tissue homogenate was prepared by using Tris-HCl buffer (0.01M) pH7.4. Serum Lipid profile (Supplementary file 2), serum LPO levels, total antioxidant capacity, OxLDL levels, and PON-1 activity were assessed in the experimental groups.

### 2.3. Methodology employed in the Present study

Total antioxidant capacity assay was assessed using a Total Antioxidant Capacity (T-AOC) Colorimetric Assay Kit (Elabscience, United States) (Catalogue No: E-BC-K219-M). The level of lipid peroxides was assayed by the method of [10]. Human serum OxLDL was evaluated using Human OxLDL ELISA Kit (Catalogue Number: E-EL-H6021) from Elabscience, United States. The levels of PON1 were measured using indirect ELISA as described by [11]. Serum PON1 activity was assessed by measuring the capacity of the enzyme to liberate p-nitro phenol based on the absorbance recorded at 412 nm during the enzymatic reaction kinetic. Blanks devoid of enzyme were used to exact for the spontaneous hydrolysis of the substrate. The expression of paraoxonase 1 (PON-1), was assessed using quantitative PCR (Primer details are provided in the supplementary file 1) and the amplified products were detected using agarose gel electrophoresis.

### 2.4. Collection of aortic tissues from patients

On arriving at certain speculations from the studies with blood samples from various risk factor as well as ischemic patients, it was further premeditated to evaluate the extent of LDL modification at the aortic tissues of patients undergoing coronary artery bypass graft surgery and compare with the serum ox-LDL levels. For comparative reasons, the aortic tissue punch samples from patients undergoing valve replacement surgery, who are generally devoid of any dyslipidemic changes, were collected. The tissues were collected after obtaining prior consent from the patients (IHEC Approval No: UM/IHEC/RM/2017-IV). The aorta tissue was fixed in 10% buffered formalin and was used for further studies.

### 2.5. Histology and Immunohistochemical analysis

Histology of aorta was studied using Hematoxylin and Eosin (H and E). The tissues were dehydrated in the descending grades of isopropanol and finally cleared in xylene. The tissues were then embedded in molten paraffin wax. Sections were cut at 5 µm thickness, stained with Hematoxylin and Eosin. The sections were then viewed under light microscope (Nikon microscope ECLIPSE E400, Japan) for histopathological changes.

Immune histochemical analysis was performed in the aortic tissue for the detection of ox-LDL, LOX-1 and PON-2 (Detailed protocol is provided in the supplementary file 1).

### 2.6. Statistical Analysis

All statistical analyses were performed using SPSS (Statistical Package for the Social Sciences) software version 20. For human studies, values are presented as n and median (Range). Based on the assessment using the Shapiro-Wilk W test, comparisons between groups were done using the Non-Parametric Kruskal-Wallis followed by Dunn’s post-hoc test and significance was determined at the level of 5% in the tables. For animal studies, values are expressed as Mean ± S.E.M (Standard Error of Mean) and statistical significance was calculated by Student-Newman-Keuls and Tukey post hoc tests, where * represents (p < 0 05), ** represents (p < 0.01) and *** (p < 0 001).

## 3. Results

### 3.1. Experiment 1 (Human Studies)

#### 3.1.1. Serum redox status, PON-1 activity and its levels

The total antioxidant capacity was found to be maximal (median being 6.1mmol/L) in the control population whereas, serum of diabetic, hypertensive as well as the ischemic patients showed significant decrement compared to control population. Despite the intra-group variations observed, intergroup comparisons also revealed decreased TAC levels in CAD patients compared to their relevant risk factor alone (Figure 1a). The levels of lipid peroxides (Figure 1b) and oxLDL (Figure 1c) were significantly elevated in all the risk factor groups when compared to healthy individuals. The level of lipid peroxides in the ischemic sub groups was found to be elevated compared to their risk factor alone counterparts. On evaluation of PON-1 protein levels (Figure 1d) and functional assessment as enzyme activity (Figure 1e), it was apparent that protein levels of PON-1 and its activity were predominantly reduced in the risk factor group when compared with that of control group individuals.

**Figure 1.**
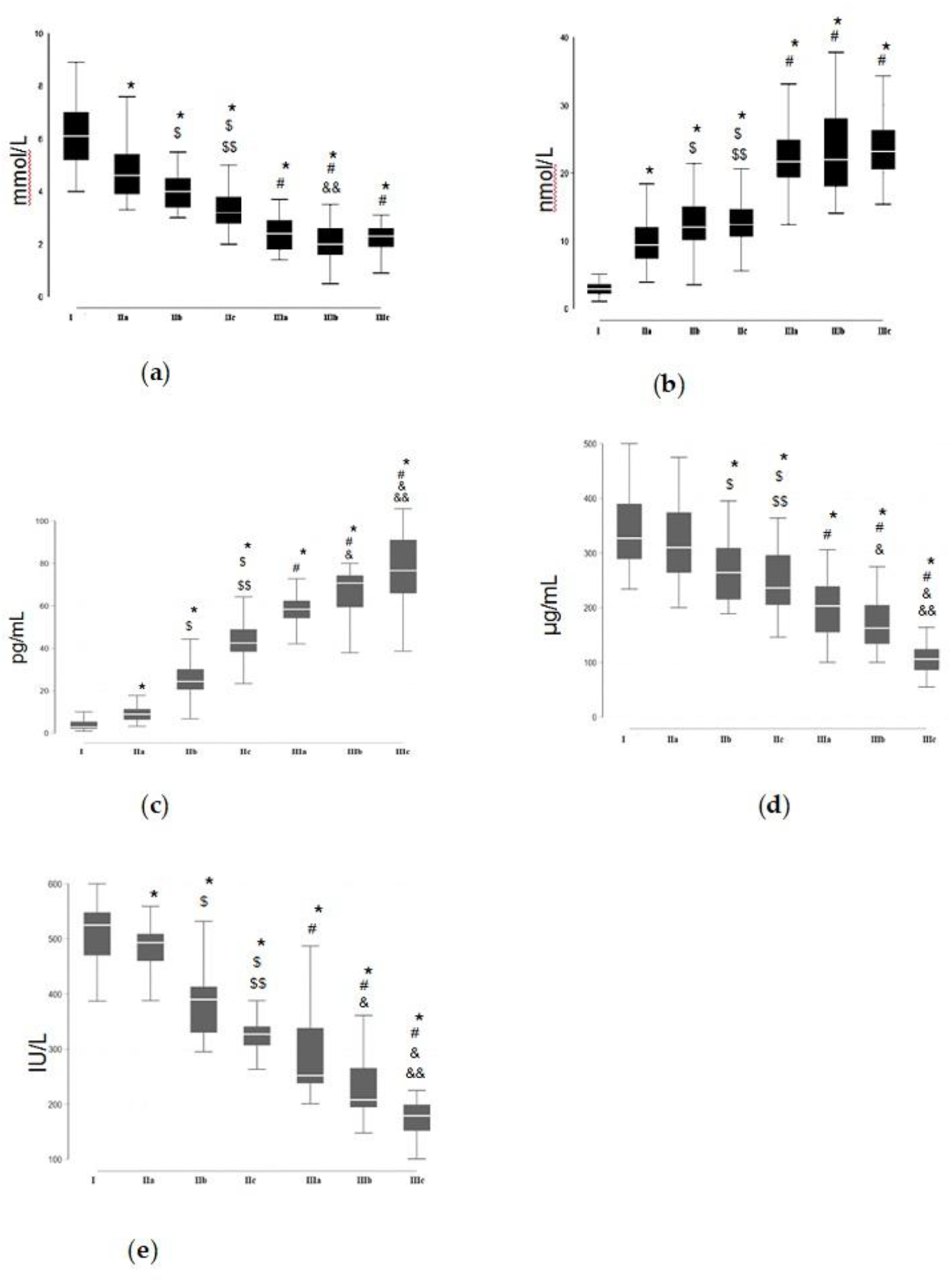
(a) Total Antioxidant Capacity; (b) LPO; (c) Ox-LDL; (d) PON-1 Levels; (e) PON-1 Activity. Values are statistically significant at the level of p<0.05 where,* represents comparison of other groups with group I controls; ‘#’ represents comparison of group III subgroups (IIIa, IIIb, and IIIc) with their respective risk factor subgroups in group II (IIa, IIb, and IIc). ‘$’ and ‘$$’ represents intra-group comparison within group II subgroups (IIa, IIb and IIc) ‘&’ and ‘&&’ represents intra-group comparison within group III subgroups (IIIa, IIIb and IIIc).

#### 3.1.2. mRNA expression of PON-1

The mRNA expression of PON-1 (Figure 2) in the aorta of CAD patients was found to be significantly lowered when compared to the control cases.PON-1 Activity decreased in the subgroups as follows: Control > HT > DM > DM+HT > CAD+HT > CAD+DM > CAD+DM+HT.

**Figure 2.**
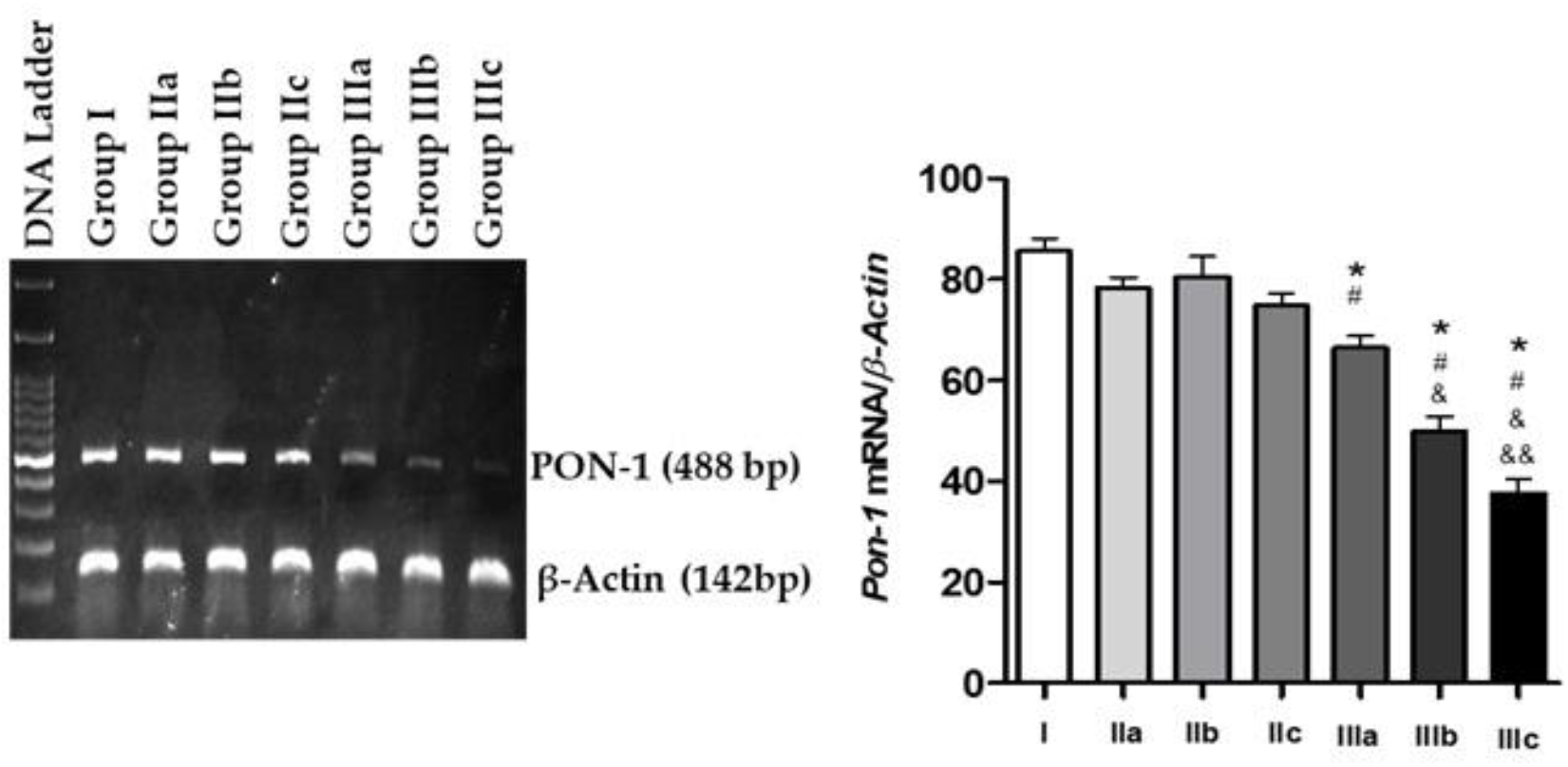
mRNA expression of PON-1 in the aorta of control subjects and patients; Values are statistically significant at the level of p<0.05 where, ‘*’ represents comparison of other groups with group I controls; ‘#’ represents comparison of group III subgroups (IIIa, IIIb, and IIIc) with their respective risk factor subgroups in group II (IIa, IIb, and IIc). ‘$’ and ‘$$’ represents intra-group comparison within group II subgroups (IIa, IIb and IIc) ‘&’ and ‘&&’ represents intra-group comparison within group III subgroups (IIIa, IIIb and IIIc).

#### 3.1.3. Histoarchitecture study of aortic punch tissues from human

Figure (3) displays histopathological sections of aortic punch of CABG patients and patients undergoing valve replacement surgery as control population using H & E staining. Control aorta displayed normal aortic tissue architecture with grooved endothelium whereas, aorta from CABG patients revealed extensive lipid accumulation in extra-cellular matrix (ECM), smooth muscle cell migration, tissue re-arrangements and neo-intimal hyperplasia along with foam cell transformation in the intimal and medial layers. Necrotic core was evidently visible and movement of smooth muscle cell towards necrotic core indicative of plaque stabilization was also noticed.

**Figure 3.**
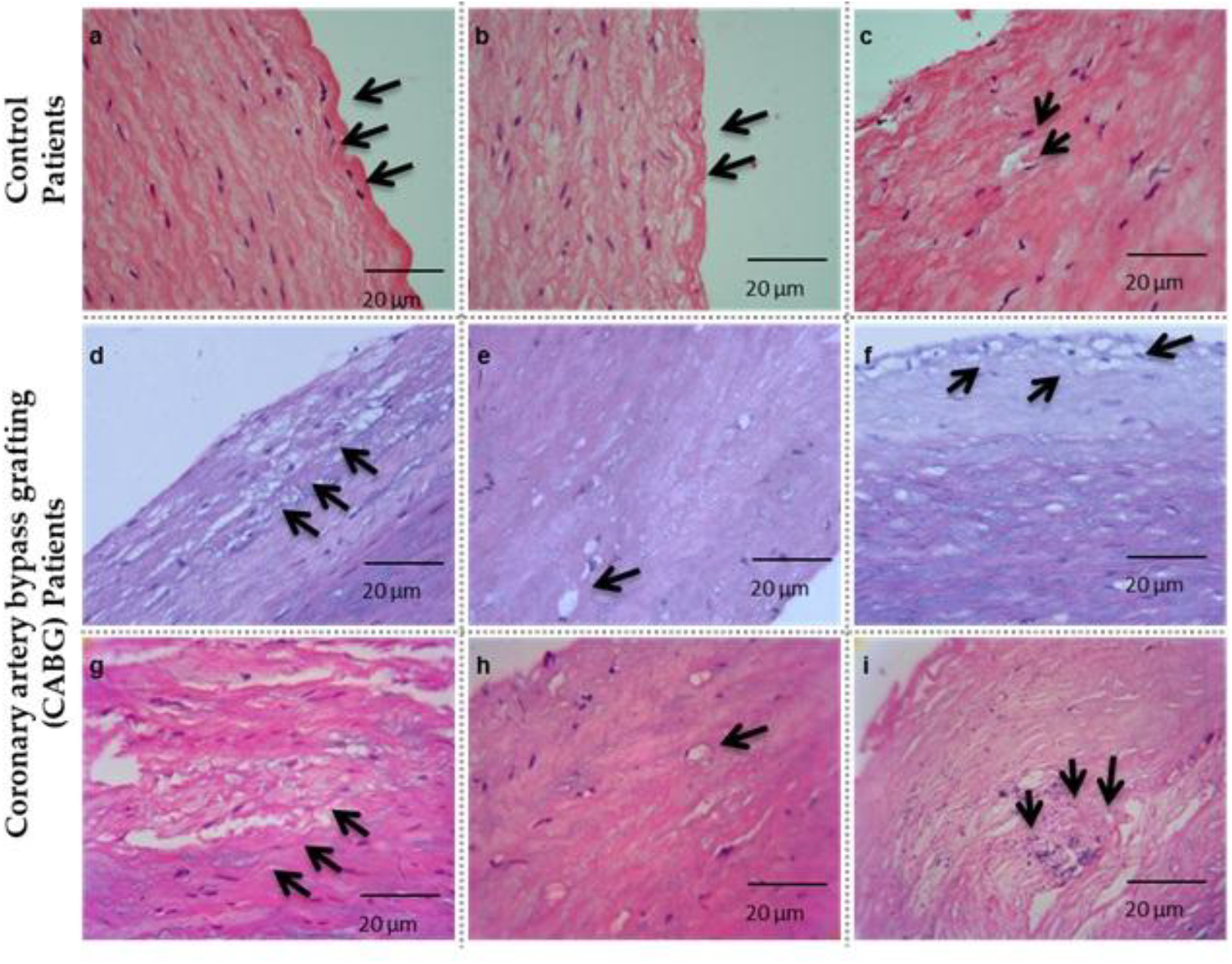
Histoarchitecture study of aortic punch tissues from CABG patients and patients undergoing valve-replacement surgery as controls. Control Aorta (a-c) shows normal tissue architecture with wavy endothelium and uniform arrangement of smooth muscle. CABG aorta (d-i) displays following characteristics: Extensive lipid accumulation and Isolated foam cell formation is seen in extra cellular matrix (ECM), smooth muscle migration, tissue re-arrangement and diffusive foam cells are evident, necrotic core formation is visible along with smooth muscle cell migration indicating plaque stabilization, extensive foam cell transformation and necrotic core destabilization are evidently visible.

#### 3.1.4. Immuno-histochemical studies of aortic tissues

Figure (4) shows the immuno-histochemical analysis of the aortic tissue samples from patients undergoing CABG as well as valve replacement surgery (as controls). Extracellular and intracellular ox-LDL have been positively stained more profoundly on the intimal layer than in the medial layer whereas, in the valve replacement patients, the ox-LDL staining was minimal or negative. The LOX-1 staining for CABG patients showed intermittent positive staining over the intimal and medial layers. On the other hand the control tissues showed clear cytoplasm and stained negative for the expression of LOX-1. The PON-2 expression was mostly negative in the cytoplasmic region of the CABG patients in both intimal and medial layers of the aortic tissues. Whereas, the aortic tissues obtained from patients with valve replacement showed higher PON-2 expression in the cytoplasm of intimal and medial layers.

**Figure 4.**
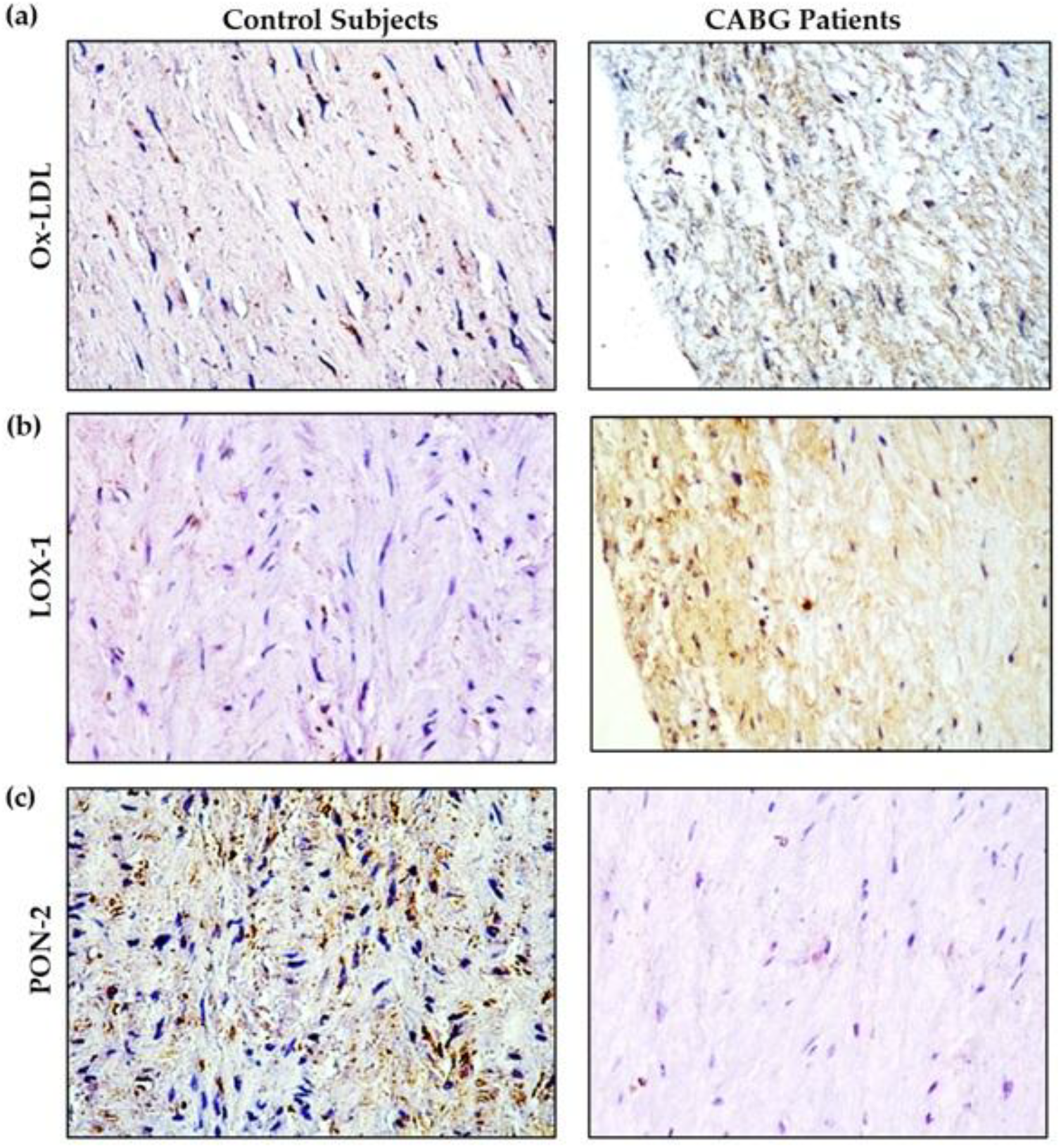
Immunohistochemical analysis of the aortic tissue samples from patients undergoing Coronary artery bypass grafting (CABG) as well as valve replacement surgery (as controls). (a) Ox-LDL - In the valve replacement patients, the ox-LDL staining was minimal or negative (left panel). Positive staining was more profound on the intimal layer than in the medial layer of CABG patients (Right panel). (b) LOX-1 - the control tissues show clear cytoplasm. On the other hand, CABG patients showed intermittent positive staining over the intimal and medial layers. (c) PON-2 - the aortic tissues obtained from patients with valve replacement show higher PON-2 expression in the cytoplasm of intimal and medial layers, whereas PON-2 expression is mostly negative in the cytoplasmic region of the CABG patients in both intimal and medial layers of the aortic tissues.

### 3.2. Experiment 2 (Animal Studies)

#### 3.2.1 Body Weight Gain

Figure (5a and b) displays the impact of high-cholesterol diet (HCD) on body weight gain compared with control rats. On feeding HCD, there was a tremendous gain in the body weight of rats fed with HCD for 45 days than that of the 15 days and 30 days.

**Figure 5.**
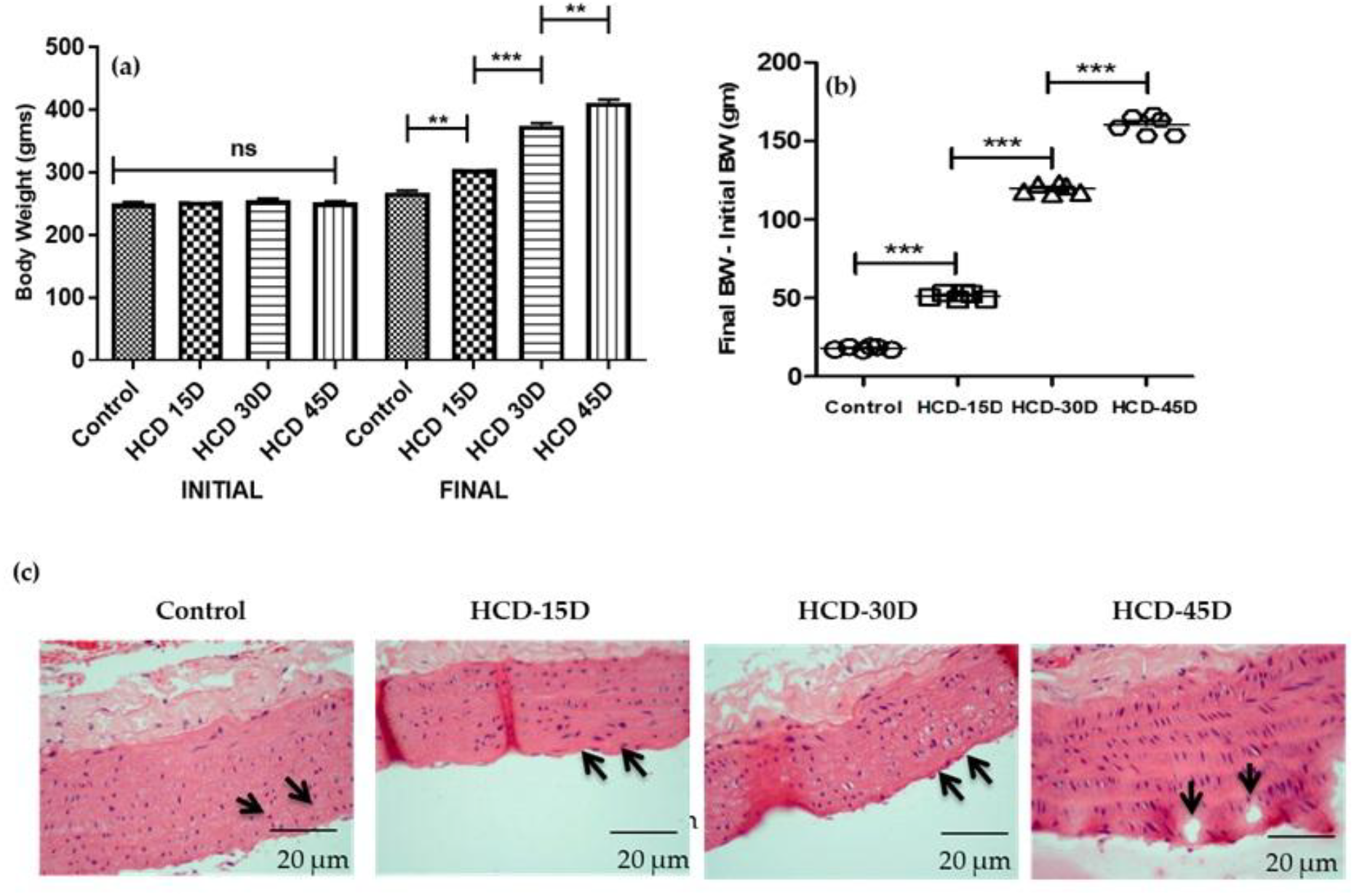
(a) Initial and Final body weight representation from control and HCD-Fed rats at different time points; (b) Body weight difference (Final – Initial). Statistical significance was calculated by Student-Newman-Keuls and Tukey post hoc tests, where * represents (p < 0 05), ** represents (p < 0.01) and *** (p < 0 001). (c) Photographic representation of the aortic wall stained with Hematoxylin and Eosin showing intimal layer thickening in the HCD-45D rats whereas such thickening is meager in the HCD-15D and HCD-30D rats.

#### 3.2.2. Histoarchitecture study of aortic tissues from rats

Figure (5c) depicts the histoarchitecture of the aortic wall from various experimental groups. Tunica intima of aorta in the control group was smooth without lipid deposition and uniformly distributed elastic fibers. Interestingly, the histopathological observations support the notion that the HCD induces the disease onset at the 30 days as there was marked increase in the perivascular deposition of fat cells and intimal layer thickening in these rat tissues whereas atherosclerotic changes is not observed in the rats fed with HCD for 15 days. Rats fed with HCD for 45 days displayed sub-intimal vacuolation, focal discontinuity of the endothelium of tunica intima with luminal adhesion of mononuclear cells to large area of denuded endothelium. Underlying tunica media shows increased thickness when compared to rats fed with HCD for 30 days.

#### 3.2.3. Serum redox status, levels of oxLDL and the activity of PON-1 in the rat serum

Extent of tissue damage was assessed by analyzing lipid peroxide levels and total antioxidant capacity in experimental animals. Assessment of MDA levels in experimental groups dictate that upon HCD feeding there is an increment in the levels of MDA and decrement in the levels of total antioxidant capacity, indicating severe oxidative imbalance (Figure 6a).

**Figure 6.**
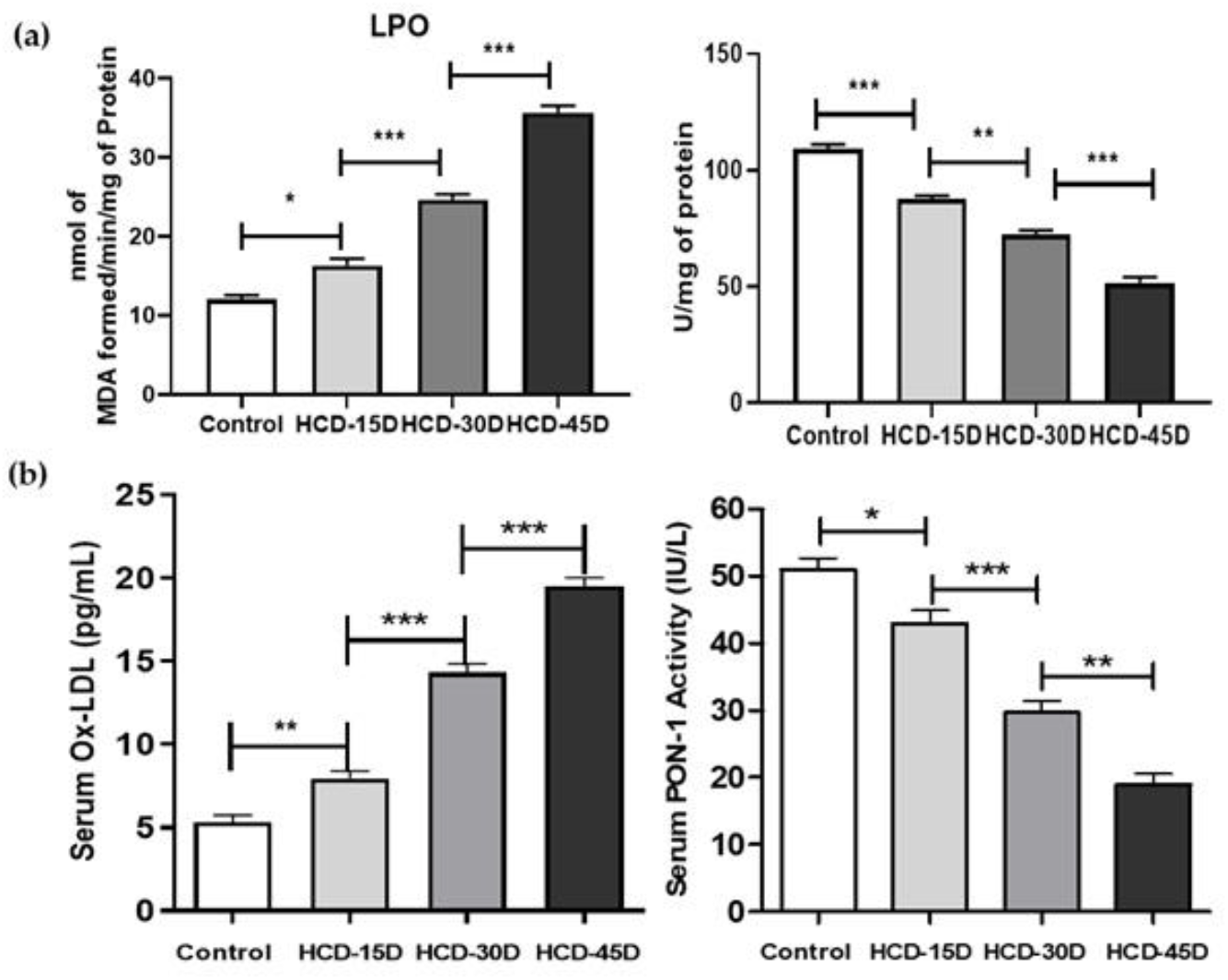
(a) Serum LPO levels and Total Antioxidant Capacity; (b) Serum Ox-LDL levels and Serum PON-1 activity from different experimental groups. Statistical significance was calculated by Student-Newman-Keuls and Tukey post hoc tests, where * represents (p < 0 05), ** represents (p < 0.01) and *** (p < 0 001).

As HCD diet altered serum lipid profile and also induces intimal layer thickening, next to determine a mechanistic link the observations was focused to determine the serum ox-LDL levels using ELISA (Figure 6b). There is a gradual elevation of serum ox-LDL in rats fed with HCD for 15 days, 30 days and 45 days. However, this elevation was significantly higher in animals administered with HCD for 45 days when compared with the rats administered with HCD for 15 and 30 days (p<0.001) which corroborates with the intimal thickening and onset of the atherosclerosis in these rats.

#### 3.2.4. Immunohistochemical analysis of the aortic tissue of the experimental animals

The immunohistochemical analysis of aorta for ox-LDL reveals increased positivity confirmed by staining of DAB in tunical layers of aorta in HCD fed rats, whereas there were no signs of positivity to ox-LDL in aorta of control animals. Assessment of ox-LDL in aorta of experimental animals dictates that ox-LDL levels elevate upon HCD feeding. 30 Days and 45 days of HCD feeding to rats showed 2 and 5 fold increase in the levels of ox-LDL when compared to control rats. While 15 days of HCD feeding doesn’t show any significant change (Figure (7).

**Figure 7.**
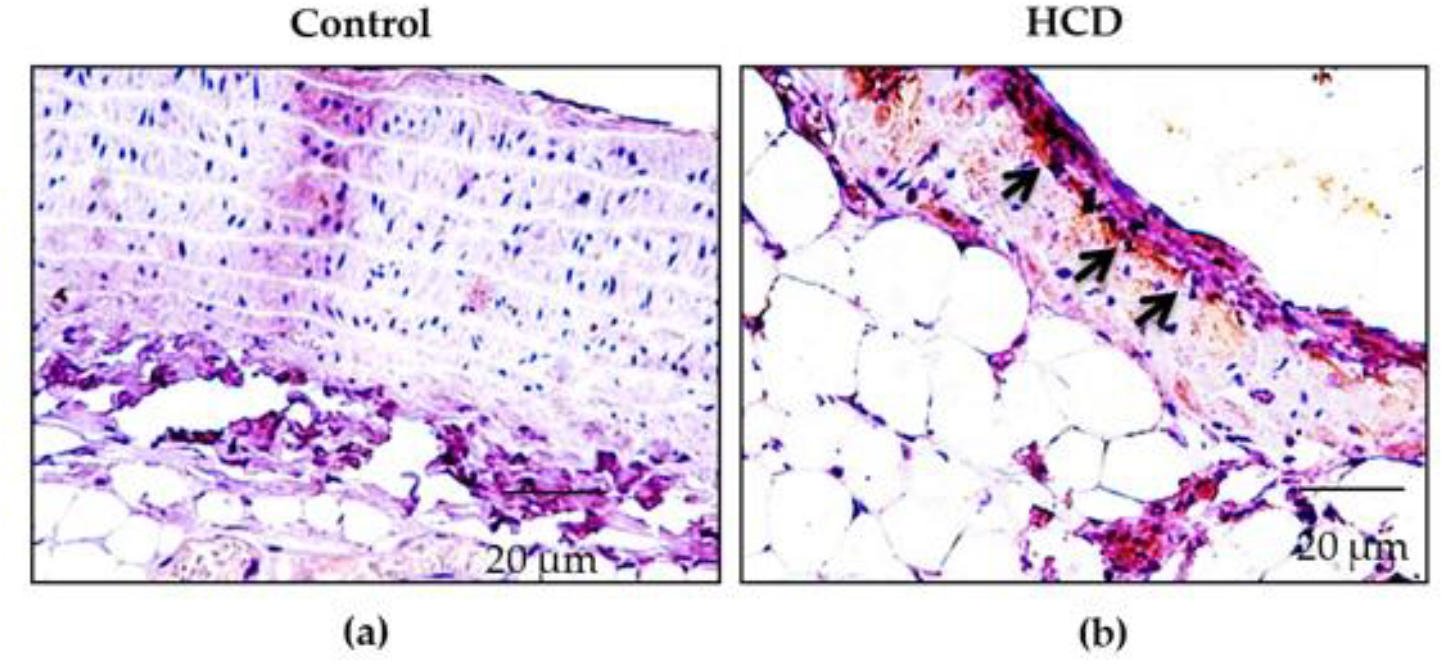
Immunohistochemical analysis of the aortic tissue of the experimental animals. (a) The control aortic tissue shows little or no expression of ox-LDL; (b) HCD fed rats show significantly higher expression of ox-LDL indicated by black arrows

## 4. Discussion

Atherosclerosis still remains a major ailment of concern in the developing countries as the consequences it imparts are quite death-defying. Diabetes and Hypertension are shown to provide favorable conditions that provoke and aggravate atherosclerosis. Despite the influential role played by these two conditions, it is undeniable that all of them do not have the tendency to develop plaques. Hence, the rationale behind devising this study is to compare and identify the attributes that discriminate the diabetic and/or hypertensive ischemic patients from their corresponding non-ischemic population who have no apparent changes.

The alteration in the lipid profiles induces endothelial dysfunction, an essential component in the pathogenesis of insulin resistance, hypertension, atherosclerosis and thrombosis [12]. In the present study, the diabetic ischemic patients with or without hypertension displayed heightened lipid anomalies and thus, hypertension and abnormal lipid profiles are known to often co-occur and coalesce to be prominent threats for CAD incidence.

Although dyslipidemic changes and accumulation of excess lipids inside the arterial walls forms one of the preliminary phases in the development of atherosclerosis, augmented intracellular reactive oxygen species act as imperative mediators of chronic inflammatory response and lead towards the progression of atherosclerosis [13, 14] by decreasing the activation of endothelial nitric oxide synthase (eNOS) and increasing ox-LDL formation [15] which necessitated to measure the levels of lipid peroxides and serum total antioxidant capacity to reflect the redox status of the system. In the present study, there is evident oxidative stress as illustrated by marked elevation of the serum lipid peroxides along with a profound decline in the serum total antioxidant capacity in both the risk factor and the ischemic groups.

According to Amponsah-Offeh et al. [16] and Griendling et al. [17], oxidative stress is relatively higher in the hypertensive patients, however, our study speculated that oxidative stress is more exacerbated in patients with a combined ailment of diabetes and hypertension (Table 3). It is undeniable that impaired redox balance accompanied with aberrations in the lipoprotein metabolism leading to the formation of ox-LDL, a major trigger in the course of atherogenesis effecting the formation of foam cells [18]. Study by Mosalmanzadeh and Pence [18] demonstrate that the oxidatively modified LDL is more culpable in the development of atherosclerosis than native LDL.

Serum ox-LDL levels are significantly elevated in both risk factor (IIa, IIb, and IIc) as well as the ischemic (IIIa, IIIb, and IIIc) subgroups of patients. Our result is in agreement with the literature which suggests the dominant role of oxidative modification of LDL in atherogenic disease progression particularly in conditions of abnormal lipid levels and disturbed antioxidative capacity [19-21]. From our results, it can be inferred that hypertension minimally modulates oxidation of LDL when compared to diabetes while, hyperglycemia co-existing with hypertension along with lipid derangements act as a dreadly triad that aggravates atherosclerotic events. Likewise, a gradual but steady increase of serum oxLDL is observed with the increase in the number of days the rats are fed with high cholesterol diet which is in harmony with the available literature [22]. Notably, there is a reduction in the activity and levels of paraoxonase, an enzyme associated with HDL is involved in protecting LDL from oxidation [23] in the cardiac patients with diabetes and/or hypertension. It is speculated that the capacity of PON1 to inactivate the oxidized lipids is attributable to the presence of an enzyme’s free sulfhydryl group at cysteine-284 [24]. Moreover, the recombinant form of PON where there was a substitution of cysteine-284 by serine is not efficient in the activation of ox-LDL [25]. This ascertains the relationship between PON and ox-LDL and emphasizes the importance to preserve PON activity during LDL oxidation. Further, mRNA expression of PON-1 isolated from peripheral blood mononuclear cells (PBMCs) shows downregulated expression of PON-1 in the ischemic subgroups hinting that parallel to the observed serum paraoxonase activity, a ubiquitous lowered expression of PON might occur in all tissues triggering atherosclerosis.

When assessing the extent of atherosclerotic changes in aortic tissues by H and E staining, it demonstrates extensive lipid accumulation in intimal and medial layers, muscle-cell migration, foam cell transformation, necrotic core formation, plaque stabilization and de-stabilization in the aortic tissue of CABG patients indicating various stages of atherosclerosis. Our reports have been supported by the earlier observations of [26, 27]. Further, histo-architecture of the rat aorta fed with HCD for 45 days shows significant atherosclerotic characteristics such as greater vacuolation, adhesion of monocyte to the tunica intima and focal discontinuity. Our results are similar to the earlier studies, where HCD intake has been shown to accelerate coronary artery atherosclerosis progression in rats [28, 29].

Further to support this, deposition of oxLDL, expression of Lectin-like oxidized low-density lipoprotein receptor-1 (LOX-1) and PON-2 were performed in the CABG patients of aortic sections by employing immunohistochemical analysis. The results reveal the presence of increased deposition of ox-LDL in atherosclerotic lesions while in the aortic tissues obtained from patients undergoing valve replacement surgery, the ox-LDL expression was found to be minimal or absent. Similar results are observed for the expression of Lectin-like oxidized low-density lipoprotein receptor-1 (LOX-1), which is mainly expressed in endothelial cells and when triggered by ox-LDL or by pro-inflammatory cytokines, up regulates the expression of pro-inflammatory signaling pathways, adhesion molecules and pro angiogenic proteins in vascular endothelial cells and macrophages [30, 31]. The expression of LOX-1 is found to be maximal in the atherosclerotic lesions whereas its expression is minimal in the aortic punch tissue of valve replacement patients. Furthermore, the expression of paraoxonase-1 which functions to scavenge the LDL from getting oxidized is observed to be reduced in the aortic punch obtained from CABG patients.

Besides, the immuno-histochemical analysis of the aorta of the rats fed with HCD reveals that the expression of ox-LDL increased significantly with the HCD feeding. The observed correlation between the ox-LDL levels and tissue damage in the present study emphasizes the elevated levels of ox-LDL in the plasma of HCD fed rats might perhaps be an indicator of the atherogenic disease progression occurring in the aorta of the rats. The Study conducted by *****et al. [32] declares that there exists equilibrium between the tissue lesions and the plasma circulating levels of ox-LDL.

## 5. Conclusions

The current study employed the assessment of atherosclerotic cardiovascular disease risk in terms of ox-LDL, which reflects the oxidative stress, parallels LDL and is a major cause for endothelial dysfunction by inducing the expression of cell adhesion molecules and PON, one of the major contributor for the atheroprotective potential of HDL.

## Ethical Approval

### Ethical Approval and Consent to Participate

All experiments were performed in accordance with the guidelines approved by the Institutional Human Ethical Committee (UM/IHEC/08-2013-I).

### Animal Studies

All experiments were perfomed as per national guidelines and protocols, approved by the Institutional Animal Ethics Committee (IAEC No:02/08/2017).

## Conflict of Interest

The authors declare that they have no potential conflict of interest to disclose.

## Acknowledgments

The financial assistance to **Prof Dr Kalaiselvi Periandavan** in the form of major grant from DST – SERB, India and to **Divya Bhavani Ravi** from University Grants Commission – Basic Scientific Research (UGC-BSR) in the form of Senior Research Fellowship (SRF), New Delhi, India is greatly acknowledged and

## Notes

### Competing Interest Statement

The authors have declared no competing interest.

## References

1. Di Cesare M, Perel P, Taylor S, Kabudula C, Bixby H, Gaziano TA, McGhie DV, Mwangi J, Pervan B, Narula J, Pineiro D. The heart of the world. Global heart. 2024 Jan 25;19(1):11.

2. La Sala L, Prattichizzo F, Ceriello A. The link between diabetes and atherosclerosis. European journal of preventive cardiology. 2019 Dec;26(2_suppl): 15–24.

3. Singh N, Shukla SK, John P, Bajpai R, Chugh P, Sengupta R, Pushkar RR, Yadav N, Singh N, Sadanandan R. Unveiling the twin epidemics of hypertension and diabetes: a cross-sectional analysis of sex-specific prevalence, risk, and hotspots in India’s epidemiological transition zones. BMC Public Health. 2025 Dec;25(1):1–21.

4. Severino P, D’Amato A, Pucci M, Infusino F, Adamo F, Birtolo LI, Netti L, Montefusco G, Chimenti C, Lavalle C, Maestrini V. Ischemic heart disease pathophysiology paradigms overview: from plaque activation to microvascular dysfunction. International journal of molecular sciences. 2020 Oct 30;21(21): 8118.

5. Poznyak AV, Sadykhov NK, Kartuesov AG, Borisov EE, Melnichenko AA, Grechko AV, Orekhov AN. Hypertension as a risk factor for atherosclerosis: Cardiovascular risk assessment. Frontiers in cardiovascular medicine. 2022 Aug 22;9:959285.

6. Poznyak AV, Nikiforov NG, Markin AM, Kashirskikh DA, Myasoedova VA, Gerasimova EV, Orekhov AN. Overview of OxLDL and its impact on cardiovascular health: focus on atherosclerosis. Frontiers in Pharmacology. 2021 Jan 11;11:613780.

7. González FE, Ponce-Ruíz N, Rojas-García AE, Bernal-Hernández YY, Mackness M, Ponce-Gallegos J, Cardoso-Saldaña G, Jorge-Galarza E, Torres-Tamayo M, Medina-Díaz IM. PON1 concentration and high-density lipoprotein characteristics as cardiovascular biomarkers. Archives of Medical Science-Atherosclerotic Diseases. 2019 Apr 12;4(1):47–54.

8. Kotur-Stevuljević J, Vekić J, Stefanović A, Zeljković A, Ninić A, Ivanišević J, Miljković M, Sopić M, Munjas J, Mihajlović M, Spasić S. Paraoxonase 1 and atherosclerosis-related diseases. Biofactors. 2020 Mar;46(2): 193–205.

9. Wamique M, Ali W, Reddy DH, Vishwakarma P, Waseem M. A case control study on HDL associated PON1 enzyme level in Northern Indian type 2 diabetes mellitus patients. Diabetes & Metabolic Syndrome: Clinical Research & Reviews. 2018 Nov 1;12(6): 843–7.

10. Senthil Kumaran, V.; Arulmathi, K.; Srividhya, R.; Kalaiselvi, P. Repletion of antioxidant status by EGCG and retardation of oxidative damage induced macromolecular anomalies in aged rats. Exp Gerontol. 2008, Mar;43(3):176–83.

11. Devasagayam, T.P.; Tarachand, U. Decreased lipid peroxidation in the rat kidney during gestation. Biochem Biophys Res Commun. 1987, May 29;145(1):134–8.

12. Hardy, F.; Djavadi-Ohaniance, L.; Goldberg, M.E. Measurement of antibody/antigen association rate constants in solution by a method based on the enzyme-linked immunosorbent assay. J Immunol Methods. 1997, Jan 15;200(1-2): 155–9.

13. Batty M, Bennett MR, Yu E. The role of oxidative stress in atherosclerosis. Cells. 2022 Nov 30;11(23):3843.

14. Masenga SK, Kabwe LS, Chakulya M, Kirabo A. Mechanisms of oxidative stress in metabolic syndrome. International journal of molecular sciences. 2023 Apr 26;24(9): 7898.

15. Yang, R.L.; Shi, Y.H.; Hao, G.; Li, W.; Le, G.W. Increasing oxidative stress with progressive hyperlipidemia in human: relation between malondialdehyde and atherogenic index. Journal of clinical biochemistry and nutrition. 2008, 43(3):154–8.

16. Amponsah-Offeh M, Diaba-Nuhoho P, Speier S, Morawietz H. Oxidative stress, antioxidants and hypertension. Antioxidants. 2023 Jan 27;12(2): 281.

17. Griendling KK, Camargo LL, Rios FJ, Alves-Lopes R, Montezano AC, Touyz RM. Oxidative stress and hypertension. Circulation research. 2021 Apr 2;128(7):993–1020.

18. Mosalmanzadeh N, Pence BD. Oxidized low-density lipoprotein and its role in immunometabolism. International Journal of Molecular Sciences. 2024 Oct 23;25(21):11386.

19. Jiang H, Zhou Y, Nabavi SM, Sahebkar A, Little PJ, Xu S, Weng J, Ge J. Mechanisms of oxidized LDL-mediated endothelial dysfunction and its consequences for the development of atherosclerosis. Frontiers in cardiovascular medicine. 2022 Jun 1;9: 925923.

20. Alique, M.; Luna, C.; Carracedo, J.; Ramírez, R.. LDL biochemical modifications: a link between atherosclerosis and aging. Food & nutrition research. 2015, Jan 1;59(1): 29240.

21. Gradinaru, D.; Borsa, C.; Ionescu, C.; Prada, G.I. Oxidized LDL and NO synthesis—biomarkers of endothelial dysfunction and ageing. Mechanisms of ageing and development. 2015, Nov 1;151:101–13.

22. Hemn, H.O.; Noordin, M.M.; Rahman, H.S.; Hazilawati, H.; Zuki, A.; Chartrand, M.S. Antihypercholesterolemic and antioxidant efficacies of zerumbone on the formation, development, and establishment of atherosclerosis in cholesterol-fed rabbits. Drug design, development and therapy. 2015, 9: 4173.

23. Durrington P, Soran H. Paraoxonase 1: evolution of the enzyme and of its role in protecting against atherosclerosis. Current Opinion in Lipidology. 2024 Aug 1;35(4): 171–8.

24. Kowalska K, Socha E, Milnerowicz H. The role of paraoxonase in cardiovascular diseases. Annals of Clinical & Laboratory Science. 2015 Mar 1;45(2): 226–33.

25. Lewoń-Mrozek D, Kurzynoga J, Jędrzejewski P, Kędzierska K, Partyka A, Kuriata-Kordek M, Ściskalska M. Molecular Structure of Paraoxonase-1 and Its Modifications in Relation to Enzyme Activity and Biological Functions— A Comprehensive Review. International Journal of Molecular Sciences. 2024 Dec 6;25(23):13129.

26. Lindeman, J.H.; Hulsbos, L.; van den Bogaerdt, A.J.; Geerts, M.; van Gool, A.J.; Hamming, J.F.; van Dijk, R.A.; Schaapherder, A.F. Qualitative evaluation of coronary atherosclerosis in a large cohort of young and middle-aged Dutch tissue donors implies that coronary thrombo-embolic manifestations are stochastic. PLoS One. 2018, Nov 27;13(11):e0207943.

27. Sonmez, F.C.; Yildiz, P.; Akhtar, M.S.; Aydin, C.; Sonmez, O.; Ay, N.; Vatankulu, M.A. NewMarkers in Atherosclerosis: Thrombospondin-2 (THBS-2) and Leukocyte Cell-Derived Chemotaxin-2 (LECT-2); An Immunohistochemical Study. Med SciMonit. 2016, Dec|31;22: 5234–5239.

28. Merchant, A.T.; Kelemen, L.E.; de Koning, L.; Lonn, E.; Vuksan, V.; Jacobs, R.; Davis, B.; Teo, K.K,; Yusuf, S.; Anand, S.S. SHARE and SHARE-AP investigators. Interrelation of saturated fat, trans fat, alcohol intake, and subclinical atherosclerosis. The American journal of clinical nutrition. 2008, Jan 1;87(1): 168–74.

29. Kamesh, V.; Sumathi, T. Antihypercholesterolemic effect of Bacopa monniera linn. on high cholesterol diet induced hypercholesterolemia in rats. Asian Pacific Journal of Tropical Medicine. 2012, Dec 1;5(12): 949–55.

30. Sánchez-León ME, Loaeza-Reyes KJ, Matias-Cervantes CA, Mayoral-Andrade G, Pérez-Campos EL, Pérez-Campos-Mayoral L, Hernández-Huerta MT, Zenteno E, Pérez-Cervera Y, Pina-Canseco S. LOX-1 in cardiovascular disease: a comprehensive molecular and clinical review. International Journal of Molecular Sciences. 2024 May 12;25(10):5276.

31. Balzan, S.; Lubrano, V. LOX-1 receptor: A potential link in atherosclerosis and cancer. Life Sci. 2018, Apr 1;198: 79–86.

32. Chen C, Khismatullin DB. Oxidized low-density lipoprotein contributes to atherogenesis via co-activation of macrophages and mast cells. PloS one. 2015 Mar 26;10(3):e0123088.

