## Supplementary 2 for "Oxidative stress and Atherosclerotic Plaque Progression: A plausible insight into the role of paraxonases and Ox-LDL"

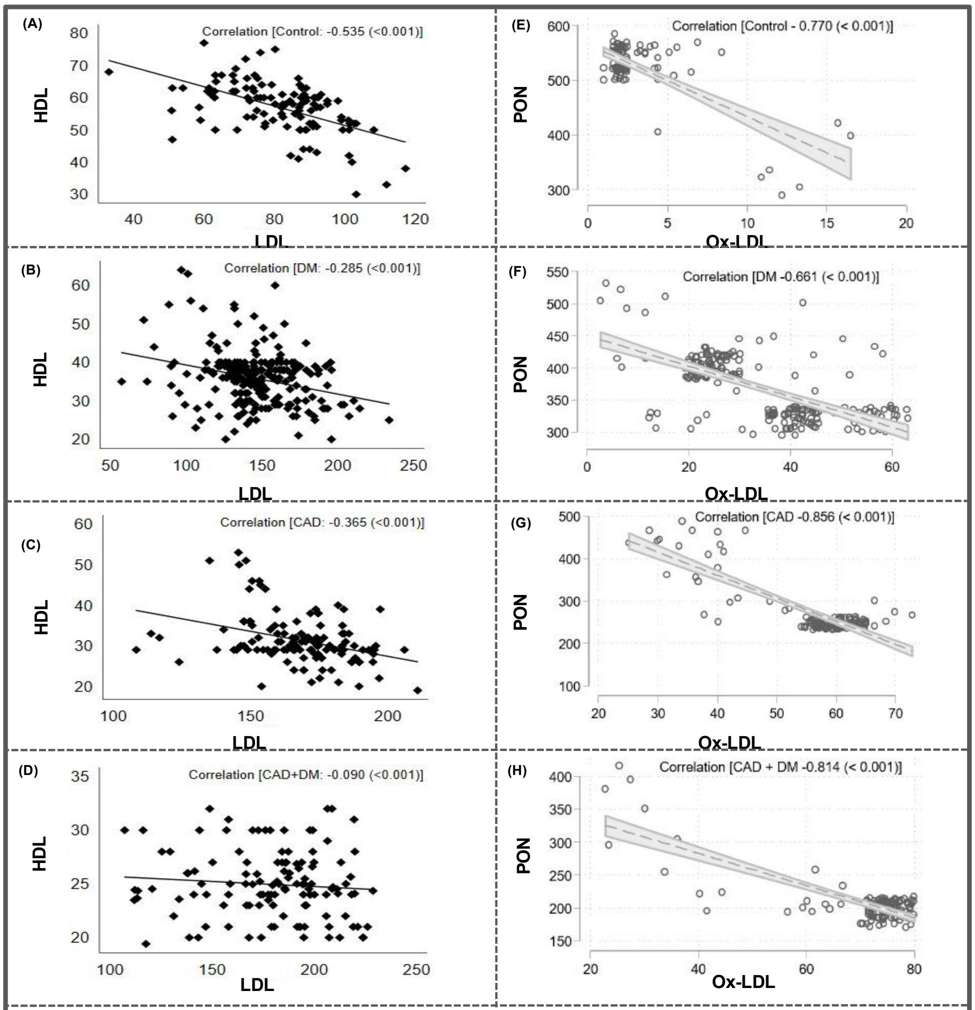


**Figure 8:** Pearson's correlation analysis of **HDL and LDL (A-D); PON and Ox-LDL (E-H)**. $p < 0.05$ significant correlation between variables. Negative RHO value in the graph indicates negative correlation between variables.

### Figure 9

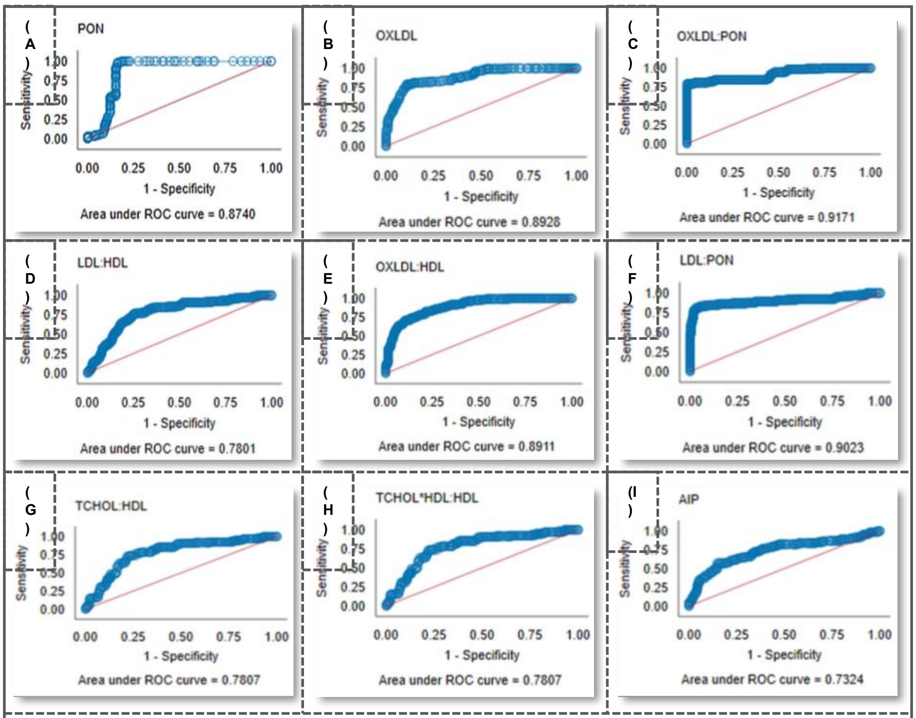


**Figure 9: Receiver-operating characteristic (ROC) curves for the biomarkers. (A) – ROC curve of PON among the study population; (B) – ROC curve of Ox-LDL among the study population; (C) – ROC curve of PON: Ox-LDL among the study population; The AUC values was appreciative for PON and Ox-LDL both independently as well in ratio combination (AUC values- 0.87; 0.89; 0.91-respectively); (D) – ROC curve of LDL: HDL among the study population. Although traditional methods substantiate the evaluation of LDL and HDL cholesterol levels for atherogenic risk assessment, the sensitivity and specificity was comparatively low for the LDL: HDL ratio; (E) – ROC curve of OX-LDL: HDL among the study population; (F) – ROC curve of LDL: PON among the study population. When the functional aspects of the lipoproteins were substituted for their levels, the resulting ratios had a significant change in the predictive values with their AUC values being 0.89 and 0.90 respectively; (G) – ROC curve of TCHOL: HDL among the study population; (H) – ROC curve of (TCHOL-HDL): HDL among the study population; (I) – ROC curve of AIP among the study population. For comparative reasons, ROC curve analysis was performed for other ratios commonly used in atherogenic risk stratifications. The AUC values of these ratios were found to be low (AUC values- 0.78; 0.78; 0.73-respectively) with a marked loss in sensitivity.**

### Table 1

| FACTORS | AUC | SENSITIVITY | SPECIFICITY | CUT-OFF |
|---|---|---|---|---|
| **PON** | 87.4% | 83.1% | 98.7% | 300 |
| **Ox – LDL** | 89.3% | 80.8% | 85.2% | 54.1 |
| **Ox - LDL : PON** | 91.7% | 80.8% | 91.3% | 0.1966 |
| **LDL : HDL** | 78.0% | 41.5% | 86.5% | 4.7655 |
| **Ox - LDL : HDL** | 89.1% | 73.1% | 86.5% | 1.3759 |
| **LDL : PON** | 90.2% | 83.9% | 92.6% | 0.5616 |
| **T.CHOL : HDL** | 78.1% | 43.1% | 87.0% | 7.0526 |
| **(T.CHOL-HDL) : HDL** | 78.1% | 43.1% | 87.0% | 6.0526 |
| **AIP** | 73.2% | 45.4% | 88.7% | 0.410 |

**Table 1: Attributes of the Receiver Operating Characteristic (ROC) analysis.**
