## Supplementary 1 for "Oxidative stress and Atherosclerotic Plaque Progression: A plausible insight into the role of paraxonases and Ox-LDL"

**Inclusion criteria**

- Individuals who consented to participate in the study (All study Groups).
- Patients with prominent risk factors like hypertension and/or diabetes (DM patients with their HbA1c levels > 6.0%; HT patients with their systolic >140 and diastolic >90) and without any apparent clinical presentation of coronary artery disease as evidenced by the patient history and recent ECG reports (Group-II).
- Patients diagnosed of coronary artery disease and are admitted in the hospital with elevated cardiac troponin levels. (Group-III).

**Exclusion criteria**

- Other Cardiac disorders
- Thyroid Disorders
- Pregnant Women
- Acute & Chronic infections
- Liver & kidney disorders
- Malignancy

**Nucleotide sequences of Forward and Reverse primer (Sigma-Aldrich) designed for RT-PCR amplification**

| **GENE NAME** | **PRIMER SEQUENCE (5’-3’)** | **ANNEALING TEMPERATURE** | **AMPLICON**  **SIZE**  **(bp)** |
| --- | --- | --- | --- |
| **PON 1**  **(forward)** | GGGAAACTGCTGATTGGCAC | 59 | 488 |
| **PON 1**  **(Reverse)** | GTCCCTGGGTGAGTACTGGA | 59 | 488 |
| **ß-ACTIN**  **(forward)** | CCGGAATGGTGAAGGTGACA | 59 | 142 |
| **ß-ACTIN**  **(reverse)** | GGGCTTCCTGAAACAATGCG | 59 | 142 |

**Immuno Histochemical Analysis**

Sections were cut at 5µm thickness using microtome and mounted on to slides and dried. Dried sections were rehydrated and heat activated antigen retrieval was done using water using pressure cooker. Tris-EDTA buffer (pH 9 (10mM Tris Base, 1mM EDTA and 0.05% Tween 20)) was added into pressure cooker and slides placed in racks, immersed in such a way that buffer covered entire section and allowed to boil for 5 minutes. Then the slides were cooled to room temperature for 30 minutes. After retrieval, sections were rinsed three times for 10 min each with 0.1 M TBS (Tris buffered saline), pH 7.4. Sections were then incubated in 1% H2O2 in TBS for 5 min and sections were permeabilized with 0.4% Triton X-100 for 30 min. The sections were blocked for non-specific binding by incubating for 30 min in 3% BSA (Bovine Serum Albumin) containing 0.2% Triton X-100 in TBS. The sections were then incubated with the primary antibody (1:100) in TBS(ox-LDL, PON-2, and LOX-1), pH 7.4 for 24 h at 4°C.After washing in TBS, sections were incubated with HRP-conjugated secondary antibody (1:1000) in TBS, pH 7.4, for 45 min at room temperature. Visualization was performed by incubation in 3,3-diaminobenzidine for 5 min. To test the specificity of the immunostaining, control sections were processed in an identical manner but with both primary and secondary antibody omitted. All sections were then washed for 10 min in TBS, mounted on slides, dried, dehydrated in increasing grades of ethanol, cleared in xylene, and mounted with DPX and cover slipped (Haycock, 1987). These sections were scanned and examined under a light microscope (Nikon Eclipse Ti series).

**The primary antibodies and their respective secondary antibody used in immunohistochemistry.**

| **S.No** | **Primary antibody** | **Source** | **Supplier** | **Dilution** | **Secondary**  **Antibody** |
| --- | --- | --- | --- | --- | --- |
| 1 | Anti ox-LDL | Mouse | Abcam | 1:100 | Donkey anti-mouse IgG-HRP conjugate |
| 2 | Anti-LOX1 | Rabbit | Abcam | 1:100 | Goat anti-rabbit IgG-HRP conjugate |
| 3 | Anti-PON2 | Rabbit | Abcam | 1:100 | Goat anti-rabbit IgG-HRP conjugate |

**Antibody and its dilution**

| S.No | Antibody | Source | M.wt | Supplier | Dilution | Secondary Antibody |
| --- | --- | --- | --- | --- | --- | --- |
| 1 | Anti-PON2 | Rabbit | 39kDa | Abcam | 1:1000 | Goat anti-rabbit IgG-HRP conjugate |
| 2 | Anti-Beta actin | Rabbit | 42kDa | Abcam | 1:1000 | Goat anti-rabbit IgG-HRP conjugate |

| **S.No** | **Primary antibody** | **Source** | **Supplier** | **Dilution** | **Secondary**  **Antibody** |
| --- | --- | --- | --- | --- | --- |
| 1 | Anti – Ox-LDL | Mouse | Abcam | 1:100 | Donkey anti-mouse IgG-HRP conjugate |
